# SCINA: Semi-Supervised Analysis of Single Cells *in silico*

**DOI:** 10.1101/559872

**Authors:** Ze Zhang, M.S. Danni Luo, Xue Zhong, Jin Huk Choi, Yuanqing Ma, Elena Mahrt, Wei Guo, Eric W Stawiski, Stacy Wang, Zora Modrusan, Somasekar Seshagiri, Payal Kapur, Xinlei Wang, Gary C. Hon, James Brugarolas, Tao Wang

**Affiliations:** Quantitative Biomedical Research Center, Department of Clinical Sciences, University of Texas Southwestern Medical Center, Dallas, TX, USA, 75390.; Bioinformatics Core Facility, University of Texas Southwestern Medical Center, Dallas, TX, USA, 75390.; Center for the Genetics of Host Defense, University of Texas Southwestern Medical Center, Dallas, TX, USA, 75390.; Kidney Cancer Program, Simmons Comprehensive Cancer Center, University of Texas Southwestern Medical Center, Dallas, TX, USA, 75390.; Department of Internal Medicine, University of Texas Southwestern Medical Center, Dallas, Texas, USA, 75390.; BioHPC, University of Texas Southwestern Medical Center, Dallas, Texas, USA, 75390.; Molecular Biology Department, Genentech, Inc., South San Francisco, CA, USA, 94080.; Bioinformatics and Computational Biology Department, Genentech, Inc., South San Francisco, CA, USA, 94080.; Department of Statistical Science, Southern Methodist University, Dallas, Texas, USA, 75275.; Department of Pathology, University of Texas Southwestern Medical Center, Dallas, TX, USA, 75390.; Laboratory of Regulatory Genomics, Cecil H. and Ida Green Center for Reproductive Biology Sciences, Division of Basic Reproductive Biology Research, Department of Obstetrics and Gynecology, University of Texas Southwestern Medical Center, Dallas, TX, USA, 75390

**Keywords:** scRNA-seq, CyTOF, SCINA, FH-deficient, RCC

## Abstract

Advances in single-cell RNA sequencing (scRNA-Seq) have allowed for comprehensive analyses of single cell data. However, current analyses of scRNA-Seq data usually start from unsupervised clustering or visualization. These methods ignore the prior knowledge of transcriptomes and of the probable structures of the data. Moreover, cell identification heavily relies on subjective and inaccurate human inspection afterwards. We reversed this paradigm and developed SCINA, a semi-supervised model, for analyses of scRNA-Seq and flow cytometry/CyTOF data, and other data of similar format, by automatically exploiting previously established gene signatures using an expectation-maximization (EM) algorithm. We applied SCINA on a wide range of datasets, and showed its accuracy, stableness and efficiency exceeded most popular unsupervised approaches. Notably, SCINA discovered an intermediate stage of oligodendrocyte from mouse brain scRNA-Seq data. SCINA also detected immune cell population shifting in *Stk4* knock-out mouse cytometry data. Finally, SCINA identified a new kidney tumor clade with similarity to FH-deficient tumors from bulk tumor data. Overall, SCINA provides both methodological advances and biological insights from perspectives different from traditional analytical methods.

## INTRODUCTION

Single cell profiling techniques such as single cell sequencing and cytometry are powerful tools for comprehensive and high-resolution characterization of cellular heterogeneities observed in tumors, brain, and other tissues. Single cell RNA-Seq (scRNA-Seq) measures the mRNA expression of several thousand genes from cells numbering a few hundred up to about 1 million, depending on the particular scRNA-Seq protocol, such as Smart-Seq (Picelli et al., 2014) or the 10X Genomics Chromium (Zheng et al., 2017). Cytometry experiments such as FACS and the recent variation, CyTOF (Cheung and Utz, 2011), can measure the expression of about 10-50 protein markers of up to 1 million cells. Many successful statistical methods, for example, Seurat (Butler et al., 2018), SINCERA (Guo et al., 2015), PhenoGraph (Levine et al., 2015) and SNN-Cliq (Xu and Su, 2015) have been developed to identify cell types from these high-dimensional profiling data with dimension reduction algorithms, unsupervised clustering and visualization techniques.

However, there are several major problems associated with unsupervised approaches. (1) Unsupervised algorithms such as K-means clustering (KC), t-SNE, *etc* only cluster the cells into groups. These cell groups are then assigned to specific cell types based on human inspection of signature genes’ expression, which is often labor-intensive and subjective, especially on borderline cases (**Fig. 1a**). Rosenberg *et al*‘s SPLiT-seq publication mentioned that they had to manually merge 73 clusters into 9 cell types *via* visual examination of the expression of cell markers (Rosenberg et al., 2018). (2) Furthermore, many cell types are identified by more than one signature genes (**Fig. 1b**). For example, CD4+ T cells need to be identified by the expression of both CD3 and CD4. These signature genes need to be manually weighed when assigning cell types, leading to even more bias and obscurity. (3) Thirdly, cell type clustering and assignment are split into two stages, where unsupervised cell clustering in the first stage of analysis ignores the knowledge of major existing cell populations and their transcriptional features. This leads to suboptimal performance especially when new cell types and subtypes are present in the sequenced cell pool. They cannot be readily differentiated in the results of unsupervised clustering. (4) Lastly, in addition to studying one experimental condition, researchers often need to assess changes between conditions in terms of the population abundances of different types of cells. Such analyses are less amenable to unsupervised approaches, as the cell groups and cell types are defined *ad hoc* each time, without justification of the reproducibility of their definitions between conditions.

**Fig. 1.**
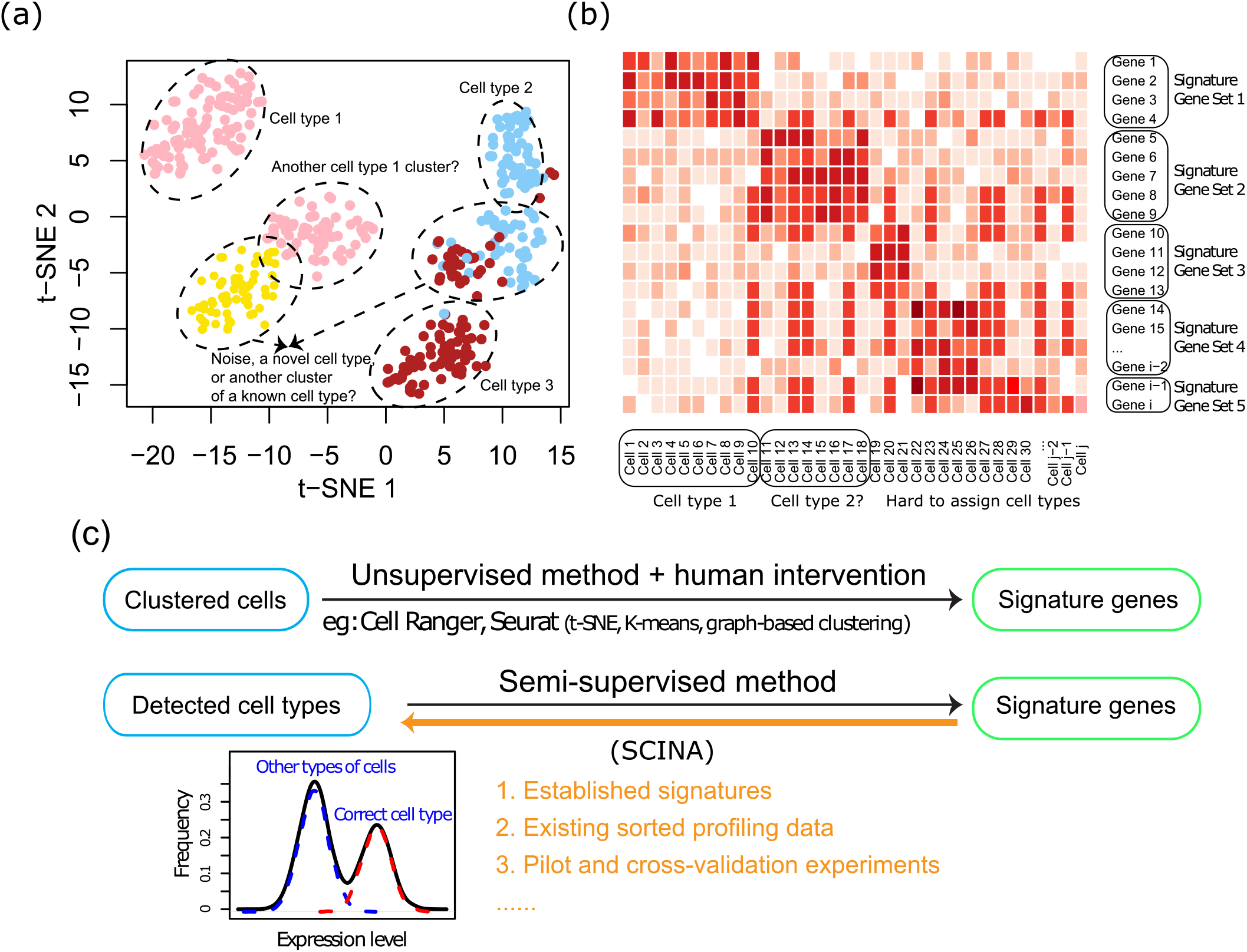
The SCINA algorithm. (a) An illustrated t-SNE plot showing the problems often associated with discerning cell types based on clustering effect of t-SNE plots. (b) An illustrated heatmap showing the difficulties of manually assigning cell types based on signature genes when multiple signature genes are known for each cell type. (c) The rationale of SCINA. Unsupervised approaches represent a one-way analysis strategy that first identifies cell categories and then relies on human knowledge for assigning cell types. SCINA represent a supervised and automated approach for assigning cell types based on bimodal distribution of expression of signature genes, which could also be applied in a reverse and interactive manner. The gene signatures could come from a variety of sources.

With the significant advancements made in high-throughput biomedical research recently, such prior knowledge of cell types and their transcriptomic features has become widely available in many cases. For example, a number of cell type-specific signature sets have been published, such as Immunome (Bindea et al., 2013) and eTME (Wang et al., 2018). Large amounts of RNA-Seq datasets of sorted cell populations are also available from databases like the Expression Atlas (https://www.ebi.ac.uk/gxa/home), which could be analyzed to define gene signatures. Alternatively, signatures could also be flexibly defined from the results of the researchers’ own pilot or crossvalidation experiments. These resources could be leveraged by an automated algorithm for both cluster detection and cell type assignment of single cell profiling data in a supervised manner.

In this study, we developed the SCINA algorithm, short for Semi-supervised Category Identification a**N**d **A**ssignment. Implemented by an expectation-maximization model, SCINA leverages prior reference information and simultaneously performs cell type clustering and assignment for known cell types (**Fig. 1c**), in a targeted manner. The prior knowledge includes a group of signature genes that are characteristically highly expressed in one type of cells, but not in other cell types. With the prior reference, SCINA searches for a segregation of the pool of profiled cells. Each subpopulation of cells highly expresses signature gene(s) specified by the researcher. The SCINA algorithm implements this by using a bio-modal distribution assumption of the expression of the signature genes. The subpopulation of cells that do not highly express the marker genes for any of the specified cell types will be designated as a novel cell type, whose exact identities can be determined in follow-up studies. SCINA is also general and can be applied to patient-level data for disease subtyping. Overall, SCINA addresses a critical research need that has been previously neglected, and proves its power for targeted cell type and subtype dissection in different scenarios of single cell profiling applications.

## RESULTS

### Validation of the SCINA model by simulation data

We first created a simulation dataset, by simulating 2000 cells of 30 cell types and one additional subset of cells as the novel cell type (**Fig. 2a**). On this dataset, SCINA yielded a classification accuracy (ACC) of 98.70%. Adjusted random index (ARI) is another metric for scoring the accuracy of categorical classifications (Zang et al., 2016), with 100% representing perfect agreement and 0% representing random guess. Our result showed a 98.46% ARI between the true labels and the SCINA classifications. On the same dataset, K-means clustering yielded an ACC of 52.70%, and an ARI of 56.97% (**Sup. Fig. 1a**), SINCERA provided an ACC of 52.72%, an ARI of 35.49%, and for PhenoGraph the ACC was 65.03% and the ARI was 43.22%. We created variations of the simulation settings, where high ACCs and ARIs for SCINA were consistently observed (**Sup. Table 1**). Next we challenged the SCINA algorithm by adding noise with four methods including: (1) “additional” gene signatures for nonexisting cell types (**Fig. 2b, top panel**); (2) simulated expression matrix with all signatures but removal of certain number of signatures from the input of SCINA (**Fig. 2b, upper middle panel**); (3) ‘redundant’ noise genes randomly selected from non-signature genes into each signature (**Fig. 2b, lower middle panel**). (4) dropouts of reads in expression values of signature genes (**Fig. 2b, bottom panel**). To our satisfaction, the ACC and ARI of SCINA remained reasonably stable across all disturbances. In particular for dropout, as SCINA relies on signature genes that are abundantly expressed, the real dropout rate is likely to be much smaller than the range covered by this simulation.

**Fig. 2.**
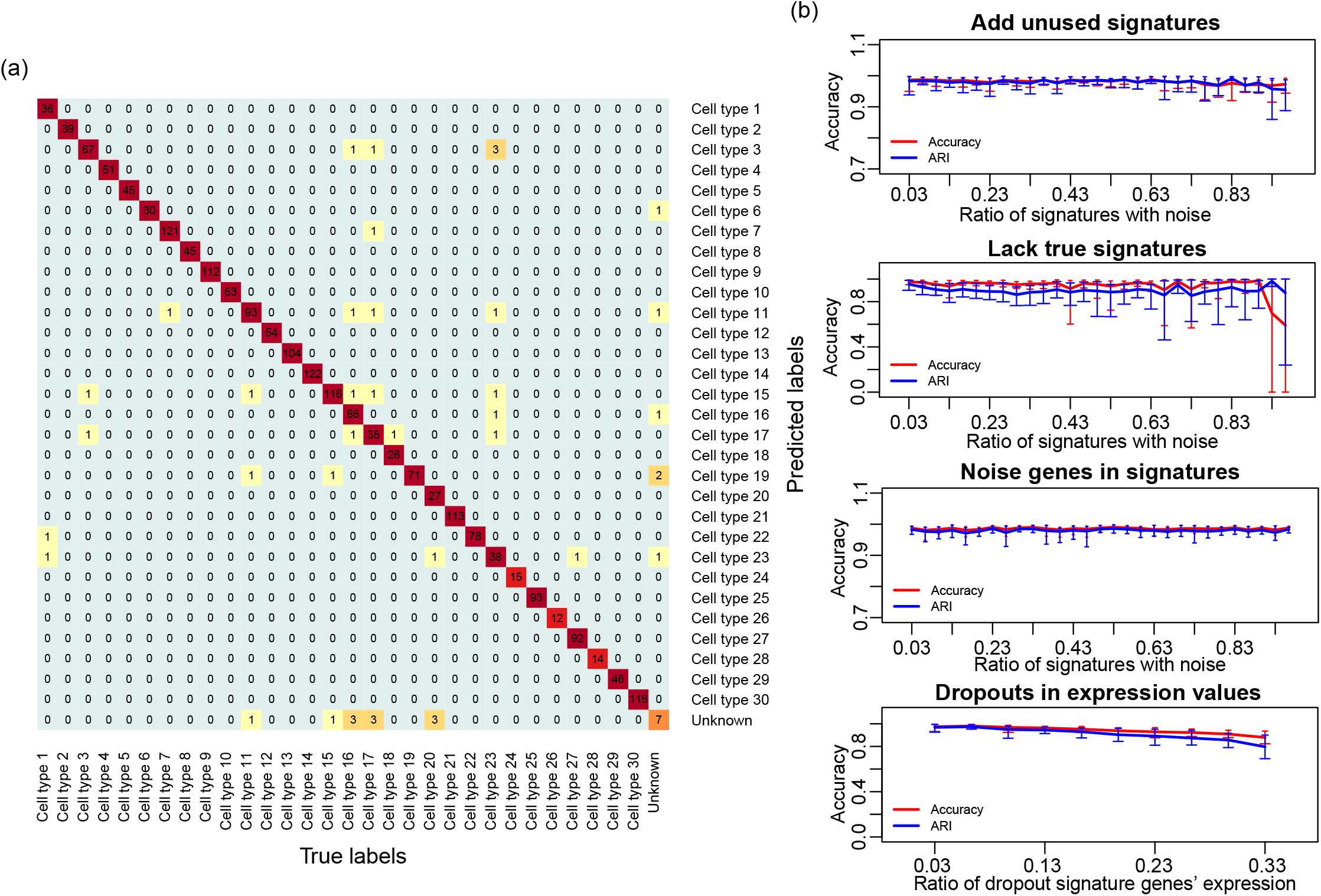
Performance of SCINA on simulated data. (a) Heatmap showing the overlap between the simulated cell types and the detected cell types by SCINA. (b) Challenging the SCINA algorithm by adding different types of noise: additional signatures for non-existent cell types, lack of true signatures for existing cell types, extra irrelevant genes in signatures, and simulated gene expression dropouts. Each scenario was repeated for 10 times, and each repeat was performed on a newly-simulated matrix of 2000 cells. Performance is judged by Adjusted Random Index (ARI) and percentage of cells correctly assigned (ACC).

We also created synthetic data based on combinations of real data, and evaluated the performance of SCINA. For this purpose, two clean populations of cells, Jurkat T cells and HEK293 (Zheng et al., 2017) were mixed. Independent expression data of these two cell lines from Dilworth *et al (Dilworth et al., 2018)* and GSE69511 were used to define signature genes. The mixing ratios ranged from 1:99 to 99:1, with a total of 2000 cells in each simulation. During cell type assignment, SCINA demonstrated accurate performance with near-perfect ARI (**Sup. Fig. 1b**) and ACC (**Sup. Fig. 1c**) across all ranges of mixing ratios. In contrast, K-means clustering (KC), SINCERA and PhenoGraph yielded unstable performance on these data, especially with unbalanced mixing ratios. Furthermore, we repeatedly simulated expression data 100 times and calculated the probability of each cell being correctly assigned to its true cell type. The average correct assignment probability of the total of 2000 cells was 98.44% for SCINA, 52.50% for KC, 52.72% for SINCERA and 18.46% for PhenoGraph (**Sup. Fig. 1d**, P<10^−5^), which indicated that SCINA achieved a much more stable performance than unsupervised clustering. Most importantly, all the unsupervised approaches we applied could not assign the clusters of cells to the exact cell types, which was done manually by us.

### Validation of the SCINA model by real data

We applied SCINA to a pool of 1,155 FACS-sorted CD45+ cells extracted from 6 clear cell Renal Cell Carcinoma (RCC) tumors (**Sup. Fig. 2**). We employed our eTME gene signatures (Wang et al., 2018) for SCINA, which are highly specific for immune cells in RCCs. In this single cell experiment, the lymphoid and myeloid cells were enriched and sequenced separately, so that the lymphoid/myeloid identity of the cells was known. We scored whether each type of detected immune cells indeed belonged to the correct sub-pool of lymphoid and myeloid cells. Dendritic cells were left out of this analysis, as they could be of either lymphoid or myeloid lineage (McLellan and Kämpgen, 2000). Overall, there was an accuracy (ACC) rate of 89.68% (**Fig. 3a**). In sharp contrast, KC yielded two clusters of cells with poor concordance with the true lymphoid/myeloid labels (ACC of 64.50%). The more advanced unsupervised methods had a moderate better performance, with SINCERA yielded an ACC of 81.03% and PhenoGraph of 69.00%. Furthermore, we conjectured that if at the pilot stage of one single cell project, the different types of cells could be sorted and sequenced to define *de novo* signatures, an even higher accuracy in subsequent experiments could be achieved. To mimic this process, we sampled 500 cells from the B cell, monocyte, and NK cell pools respectively from Zheng *et al* and defined a set of *de novo* gene signatures (Zheng et al., 2017). Then the rest of these types of cells (n=29,345) from the same study were mixed, along with CD4 T helper cells as the pseudo “unknown” cell type. As we expected, SCINA achieved a 97.3% ACC on this dataset (**Fig. 3b**). Meanwhile, KC yielded an ACC of only 50.2%, Seurat provided a prediction ACC of 64.44%. The mixed dataset exceeded the optimized data scale of SINCERA and PhenoGraph, so we randomly select a subset containing 10% of the cells. The ACC performance of SINCERA and PhenoGraph on the subset was 54.44%, and 56.26%. Next, we downloaded the CyTOF data from Hawley *et al* (Hawley et al., 2017). In this study, the mouse lacrimal gland was acutely injured *via* intraglandular injection of IL-1α, and the lacrimal gland single cell suspensions from day 1 to day 7 following injury were collected and subjected to CyTOF analysis. SCINA was applied on this dataset with established protein surface markers and discovered that neutrophil contents peaked at day 1 following injury and monocyte contents peaked at days 2 and 3 (**Fig. 3c**), which is consistent with the overall conclusion of the original publication. In comparison, t-SNE was applied to cluster and visualize the single cells. The t-SNE plot (**Fig. 3d**) shows that cells of the same type, assigned by SCINA, indeed cluster together, confirming the accurate performance of SCINA. However, there is a lack of clear boundaries between different types of cells on the t-SNE plane, making it impossible to use t-SNE as the sole methodology for assigning cell types in CyTOF data.

**Fig. 3.**
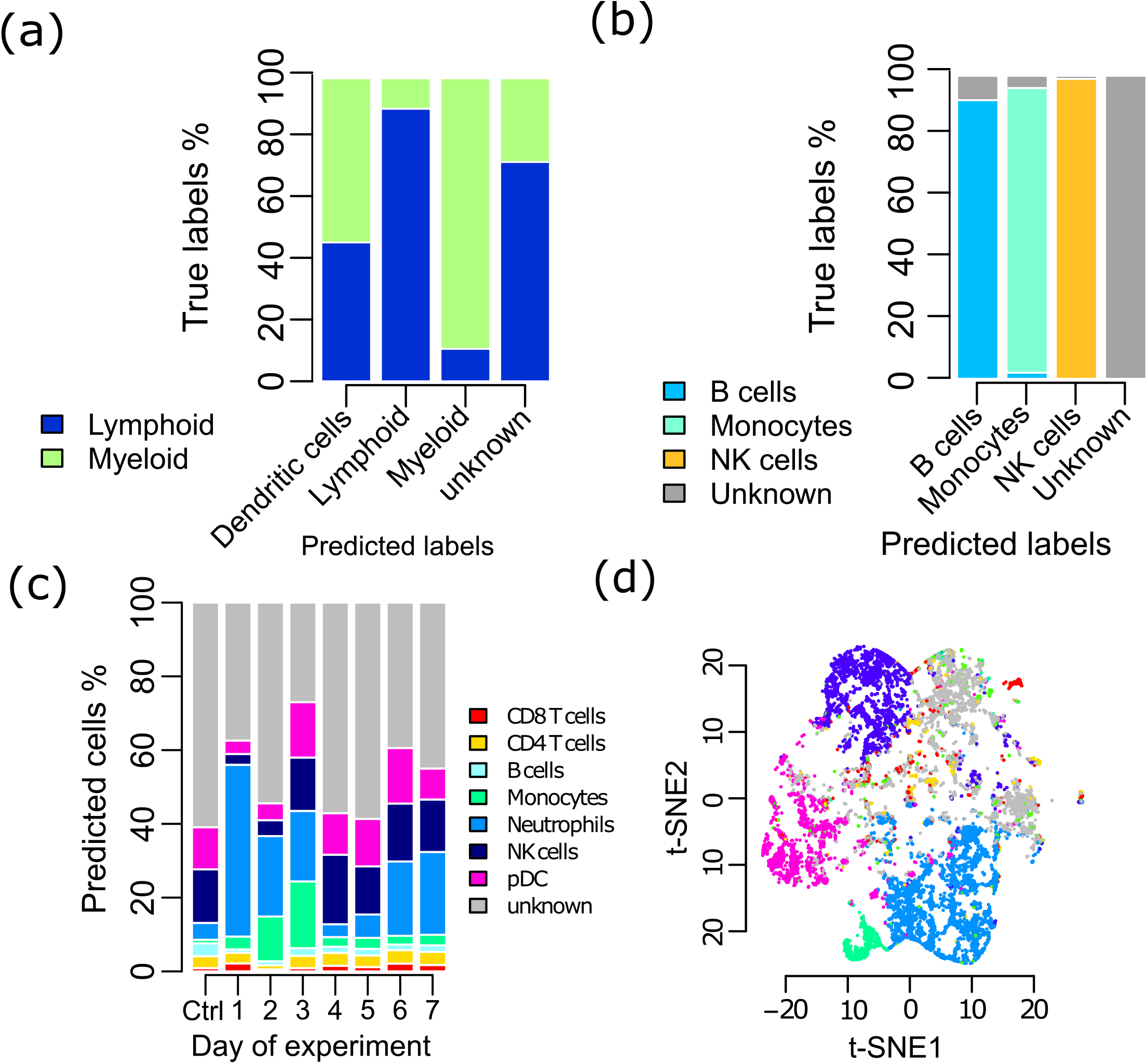
Validation of SCINA on real data (a) SCINA identified the cell types of CD45+ single cells enriched from RCCs. Dendritic cells were left out of this analysis, as they could be of either lymphoid or myeloid lineage. (b) SCINA identified the cell types in a pool of cells comprised of B cells, monocytes, NK cells, and a “pseudo” unknown cell type. (c) SCINA was used to analyze the mouse CyTOF data collected each day following gland injury, which profiled an average of 389,777 cells at each time point. (d) t-SNE was used to analyze the same mouse CyTOF dataset. The cells were colored by cell types assigned by SCINA

### Discovery of a new stage of oligodendrocyte development in mouse brain

To demonstrate the capability of SCINA to derive novel biological discoveries, we applied SCINA on the mouse brain scRNA-Seq data from Rosenberg *et al* (Rosenberg et al., 2018). The original publication performed cell type assignment based on painstaking manual inspection of unsupervised clustering results. In contrast, SCINA automatically generated cell type predictions for the 27,096 nonneuronal cells with the same markers used by Rosenberg *et al* but slightly expanded to also include the most positively correlated genes. Overall, the SCINA-assigned cell types for most cells were consistent with those by the manual inspection method (**Fig. 4a**). However, the manual method leads to the intertwined boundaries of the oligodendrocytes and OPC cells (the circles in **Fig. 4a**). In comparison, SCINA divided the cells in this region into oligodendrocytes, OPCs and an unknown cell type. The density plots of cell type assignment probabilities (**Fig. 4b**) demonstrate confident and clear separations between these three types of cells. We examined the expression of*Mbp*, marker for oligodendrocytes; and *Pdgfra*, marker for OPCs, which were used by Rosenberg *et al.* The findings also supported the grouping of these cells into an independent type (**Fig. 4c**). In fact, these cells indeed seem to possess a unique transcriptomic program. We identified genes that were up-regulated in the unknown cells but down-regulated in both oligodendrocytes and OPCs. One of the top genes was *Tmem108*, which has been previously linked with schizophrenia and alcoholism (Heath et al., 2011; O’Donovan et al., 2008) (p-values<10^−5^ for comparing these cells with OPCs and oligodendrocytes, **Fig. 4d**). Gene ontology analysis also confirmed the existence of differentially enriched pathways related to cell projection regulation in these cells (**Sup. Table 2**). Overall, the unknown cell type marked by SCINA could likely represent a newly-defined intermediate stage between OPC and oligodendrocyte development in mouse.

**Fig. 4.**
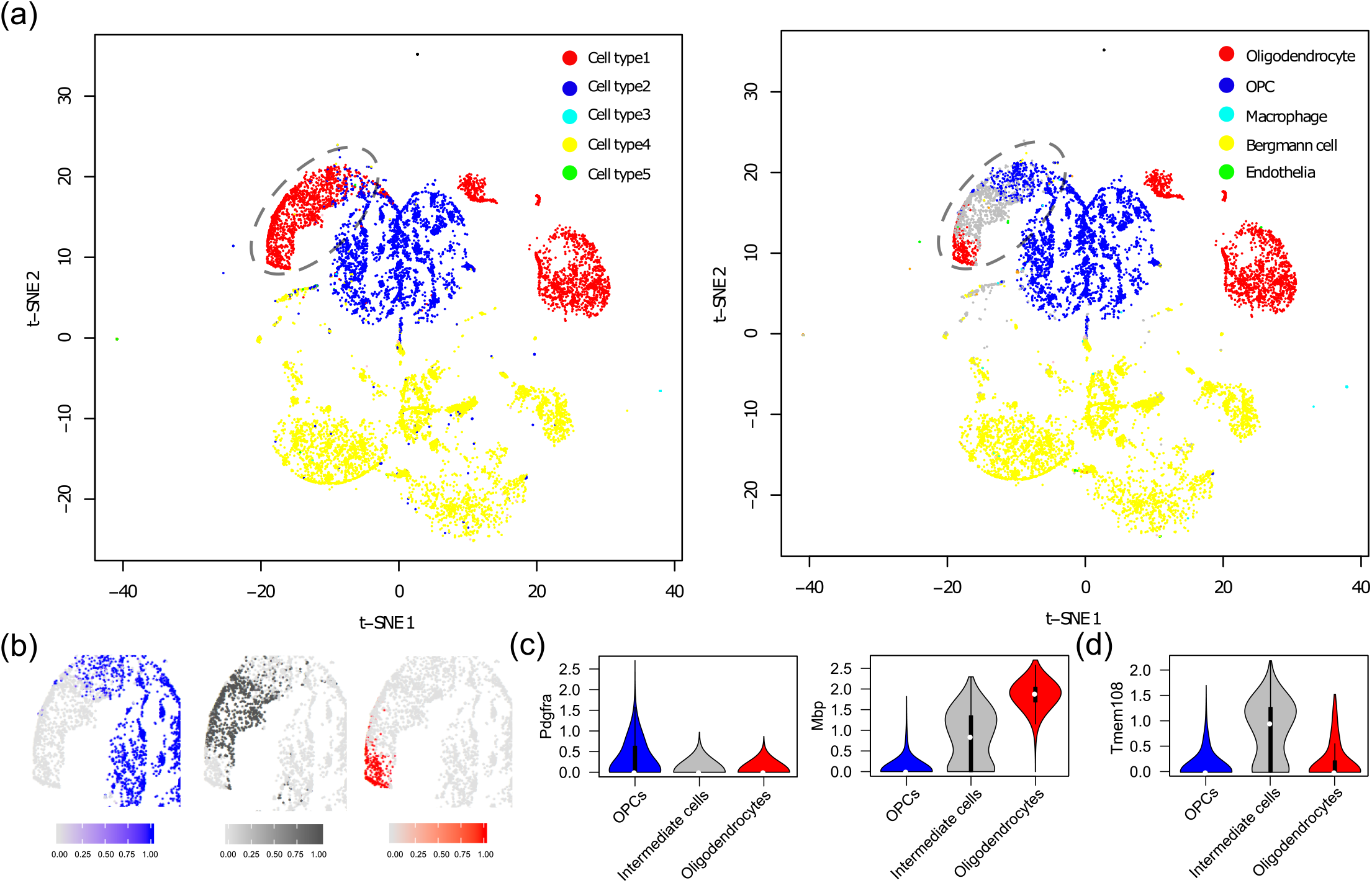
Discovery of a new stage of oligodendrocyte development in mouse brain. (a) t-SNE plot showing the clusters of cells detected by the manual inspection method employed in the original publication (left) and the cell types assigned by SCINA (right). (b) Density plots of cell type assignment probabilities generated by SCINA. (c) Violin plots showing the expression of Mbp and Pdgfra and (d) violin plot showing the expression of Tmem108 in OPCs, oligodendrocytes, and the intermediate stage of cells.

### SCINA detected immune cell population alterations in *Stk4* knock-out mice

Next, we evaluated SCINA on cytometry data. We generated a pedigree of CRISPR knock-out (KO) mice in *Stk4*, which is a key regulator of the Hippo pathway (Zhao et al., 2010). The blood sample of each mouse was run through a standard FACS pipeline as previously reported (Wang et al., 2015). We had established a manual serial gating schema to sort and identify the populations of each type of immune cell from the FACS data using FlowJo (https://www.flowjo.com/). These FACS data were also analyzed by SCINA based on commonly used cell surface markers. We investigated the abundances of T cells, B cells, NK cells, macrophages, and neutrophils from all mice in the context of the *Stk4* genotype. Both the SCINA and manual gating methods detected lower T cell levels (P<10^−5^ for both SCINA and serial gating), and elevated B cell levels (P=3.93×10^−4^ for SCINA, P=1.87×10^−3^ for serial gating) and NK cell levels (P=0.008 for SCINA, P<10^−5^ for serial gating) (**Fig. 5a**). The T cell phenotype is consistent with previous reports in both human and mouse models that observed a correlation between *Stk4* deficiency and a paucity in T cells (Abdollahpour et al., 2012; Bai et al., 2016). However, the manual gating method concluded that *Stk4* KO led to a significant increase in both neutrophils (**Fig. 5b**, P<10^−5^) and monocytes (**Fig. 5c**, P<10^−5^), while the neutrophil and monocyte counts are not significantly different between wild type and KO mice according to SCINA (P=0.976 and P=0.577). To resolve this conflict, we carried out a Complete Blood Count (CBC) analysis using blood from this pedigree of mice. We confirmed the insignificant changes of neutrophil and monocyte counts after knocking out *Stk4* (P=0.496 for neutrophils and P=0.209 for monocytes), which were in concordance with the SCINA results. Overall, these experiments demonstrated the ability of SCINA to yield biological discoveries while pruning spurious errors of manual cytometry analytical methods.

**Fig. 5.**
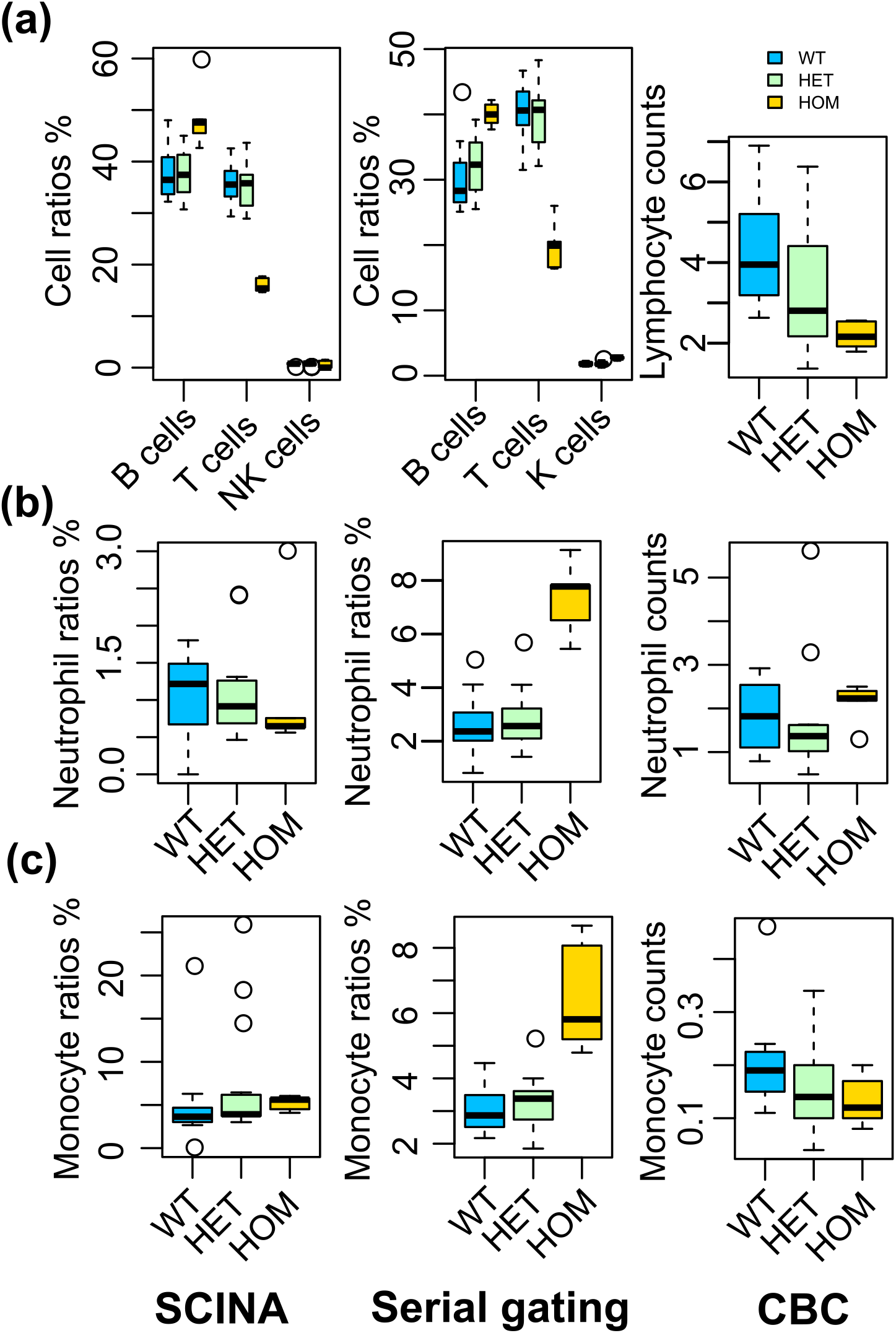
SCINA detects immune cell alterations in Stk4 KO mice. The pedigree contained a total of 12 wild-type (WT) mice, 15 heterozygous (HET) mice, and 5 homozygous (HOM) mice. The relative levels of immune cell populations are detected by SCINA from FACS data (left), the serial gating method from FACS data (middle), and CBC (right). (a) Lymphocytes, (b) Neutrophils, and (c) Macrophages.

### SCINA identified a novel tumor clade based on patient bulk RNA-seq profiles

SCINA also solves a general supervised classification problem, for example, for patient-level sequencing data. RCCs are mainly comprised of clear cell RCC (ccRCC), papillary RCC (pRCC), and chromophobe RCC (chRCC). One minor but aggressive subtype of RCC has been recently identified: HLRCC, which bears morphological similarities with high-grade pRCCs and is characterized by germline mutations in the Fumarate Hydratase (*FH*) gene (Smith et al., 2016). We collected TCGA pan-RCC RNA-seq data from the Broad GDAC Firehose and also from the Kidney Cancer Program (KCP) at UT Southwestern Medical Center(Durinck et al., 2015). Preliminary pathological reviews determined that these RCCs are comprised of 528 ccRCCs, 323 pRCCs, 45 chRCCs, and 11 FH-deficient (FHD) samples (9 with germline mutations and 2 with somatic mutations in *FH*), along with 163 adjacent normal kidney samples (**Table 1**). Assuming most of these pathological classifications were correct, we defined gene signatures for each tumor type and for the normal kidney tissue based on differential expression analyses. SCINA was then applied with these gene signatures on the transcriptomic data to re-classify the patient subtypes, which resulted in a 91.9% concordance rate when comparing with the original pathological review (**Table 1**). However, there are 16 TCGA ccRCCs by pathological standards that were reclassified as chRCCs by SCINA. Interestingly, a recent second pathological review of the TCGA cohort suggested that 11 of these 16 RCCs could indeed be chRCCs (Ricketts et al., 2018).

**Table 1.**
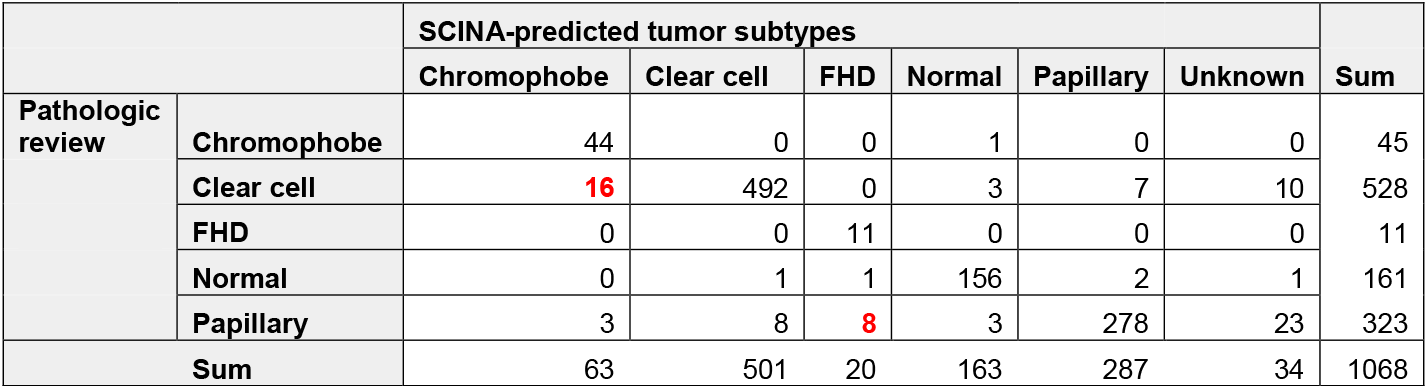
Overlapping RCC subtypes assigned by pathological reviews and subtypes assigned by SCINA.

Most interestingly, SCINA classified 8 pRCCs tumors as FHDs, which inspired us to carry out in-depth analyses to resolve this conflict. First, Principal Component Analysis (PCA) based on the whole transcriptome (**Fig. 6a**) confirmed that these 8 tumors clustered closely with the 11 FHDs. Pathological review by a GU pathologist revealed morphology features indistinguishable between FHDs and these 8 tumors (**Fig. 6b**). However, these 8 tumors did not have any *FH* mutations (**Fig. 6c**). Thus, they are not FHDs by definition, but could represent a closely related tumor clade discovered by SCINA analysis of RNA-Seq data. We named them “FHD-like” and examined whether FHD and FHD-like tumors share convergent molecular perturbations. Interestingly, both FHDs and FHD-like tumors have enriched mutations in the Hippo pathway *(NF2* and *PTPN14)* and recurrent loss of p16 expression (in-house cohort) (**Fig. 6c**), which are well-characterized tumor suppressor genes (Liggett and Sidransky, 1998; Sebio and Lenz, 2015). The TCGA analysis of the pRCC data demonstrated hypermethylation of *CDKN2A* (p16) in the 5 TCGA FHD-like tumors (Cancer Genome Atlas Research Network et al., 2016).

**Fig. 6.**
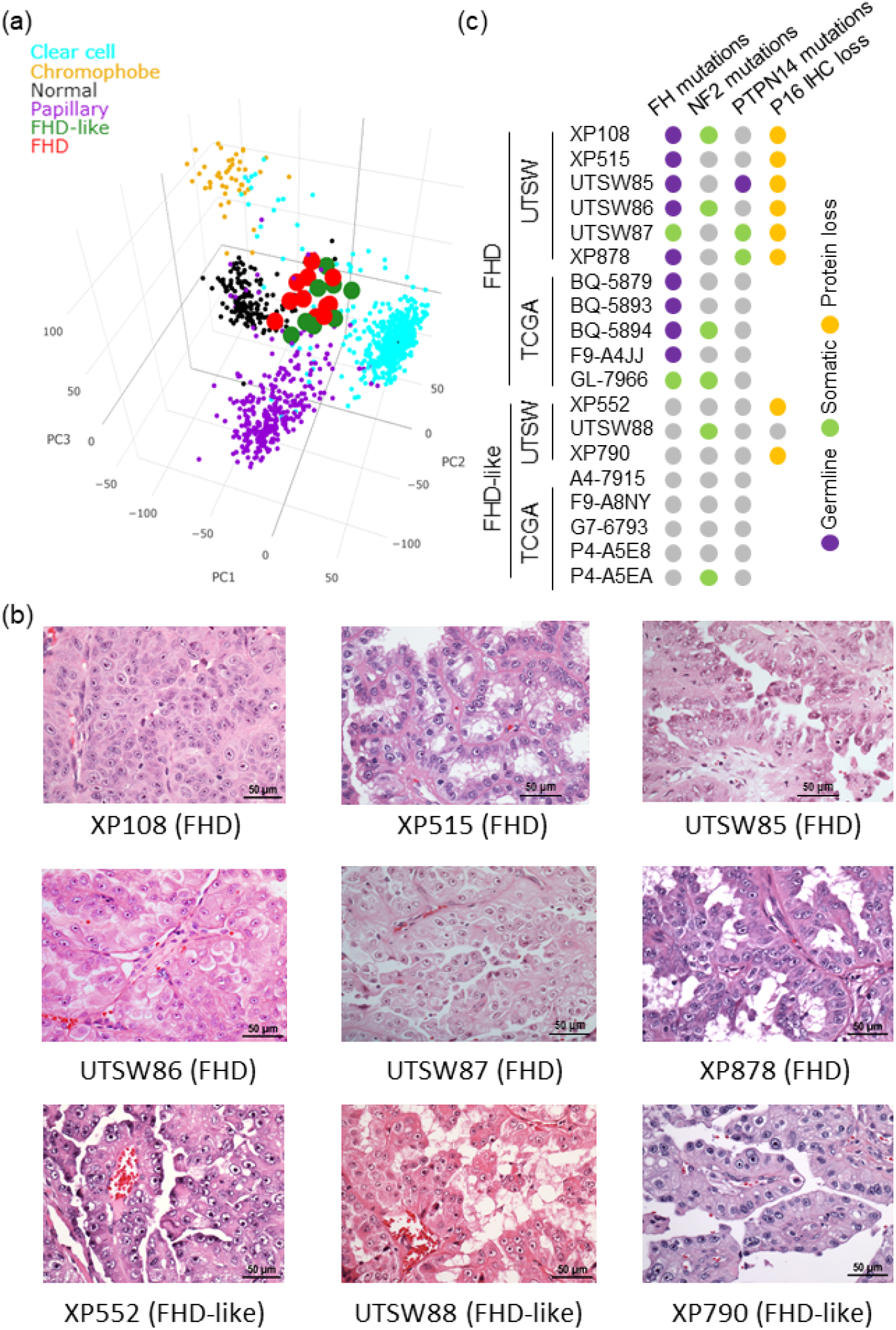
SCINA identified a novel RCC tumor clade based on gene expression profiling. (a) 3d PCA plot showing clustering of UTSW KCP and TCGA RCC and normal kidney samples by expression (n=1,068). (b) H&E stained sections showing that both FHD and FHD-like tumors had features characteristic of HLRCC, including papillary architecture, large nuclei, and prominent eosinophilic macronucleoli, with perinucleolar clearing. (c) Somatic/germline mutations and IHC results for FHD and FHD-like tumors.

Overall, these analyses attest to the capability of SCINA to correct the mis-classifications of human pathologists and to discover new entities of patient subtypes.

### The SCINA R package and SCINA on the cloud

For convenience of biologists and bioinformaticians who wish to apply SCINA in their research, we have created an R package (https://github.com/jcao89757/SCINA and https://cran.r-project.org/web/packages/SCINA) and also a web server (**Fig. 7a**, http://lce.biohpc.swmed.edu/scina) to host SCINA on the cloud. Importantly, the R package/webserver has provided a functionality (the heatmap) for users to visually assess the degree of bi-modal distribution of each signature gene in the particular application. SCINA is extremely computationally efficient. On the same datasets, SCINA is 20 times to over 100 times faster than Seurat, t-SNE, and Cell Ranger, and 200 to 400 times faster than SINCERA and PhenoGraph (**Fig. 7b**).

**Fig. 7.**
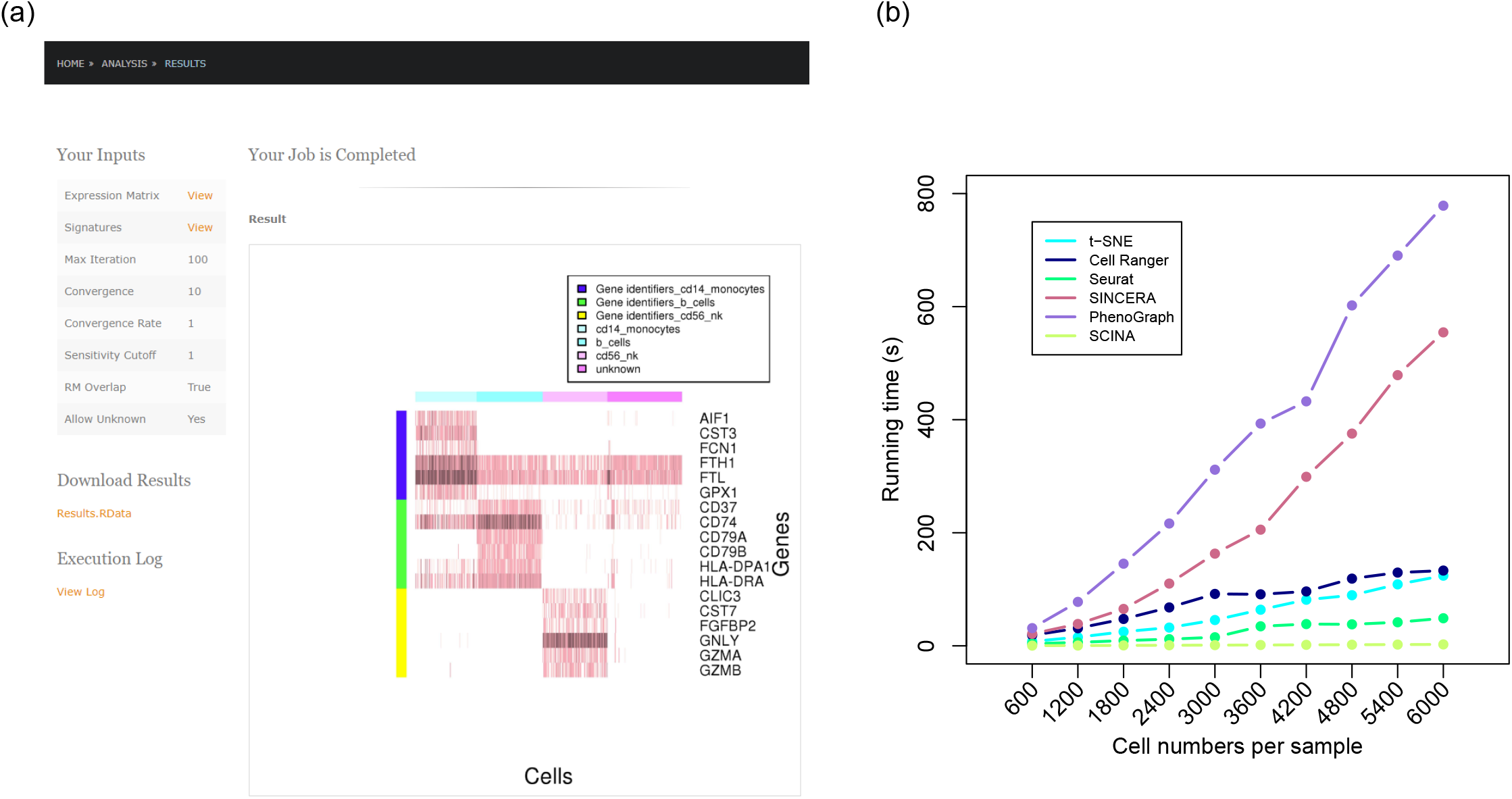
The SCINA R package and web server. (a) Screenshot of the SCINA web server. (b) Runtime comparison for SCINA, Cell Ranger (K-means clustering mode with graph-based fine-tuning), t-SNE, Seurat, SINCERA, and PhenoGraph. Runtime tests were performed on 10 datasets, constructed with 280n randomly selected HEK293 cells (out of 2,885 total cells) and 320n randomly selected Jurkat cells out of 3,258 total cells), where n was an integer ranging from 1 to 10. Both HEK293 cell and Jurkat cell sets were publically available from Zheng et al.

## DISCUSSION

SCINA is a novel methodology for cell type classifications in scRNA-Seq or CyTOF/FACS data. Compared with unsupervised approaches, SCINA accepts the signatures of any number of marker genes, and weighing of the genes for each cell type is determined automatically by the algorithm. It is also generally applicable as a classification algorithm when data of similar formats are available. In our study, the performance of SCINA was comprehensively validated on a variety of datasets, which showed the accurate performance of SCINA and its superiority to other unsupervised software. This superiority is likely to be more pronounced when the data analysis problem is more difficult, such as the difficult problem of dealing with 30 cell types in **Fig. 2** and the unbalanced mixing proportions in **Sup. Fig. 1bc**. SCINA requires less manual intervention and is more robust and objective. Despite built upon the bi-modality assumption, SCINA does not necessarily require each gene’s expression to have totally clear and wide separation of the two modes. Some of the signature genes may have gradual changes along a differential or functional trajectory. Instead, SCINA relies on the separation of clusters of cells in a high-dimensional space formed by the contribution from all the signatures genes.

Nevertheless, SCINA is synergistic and complementary to these other trending methods. For example, the unsupervised approaches can be more appropriate for the data exploration stage, while SCINA’s semi-supervised model provides more consistent classification of cells when preliminary knowledge has been gained. By employing them in an interactive manner, one project may start by applying unsupervised methods like t-SNE and *ad hoc* analyses for visualization and to identify new cell types or subtypes. The researchers may then use SCINA and the *de novo* signatures to detect the existence of these newly defined cell populations in subsequent experiments to understand how they change under different perturbation conditions.

Prior knowledge of signature genes is available to the researchers in many biomedical research settings, which could come from several sources (**Fig. 1c**): (1) previously published signatures, (2) pre-existing sequencing data of sorted cells, and (3) pilot or cross-validation experiments where *de novo* signatures may be defined. (3) is a completely novel but very useful application of SCINA that cannot be afforded by unsupervised methods, which allows the researchers to assess the reproducibility of their biological phenomenon of interest across experimental conditions and replicates. SCINA offers the flexibility of using any signature set defined by researchers that is tailored for the biological problem under investigation. On the other hand, SCINA could also be regarded as a discovery tool for novel cell type-specific signatures (**Fig. 1c**). For example, cell types in the single cell profiling data can be defined using a smaller pre-defined set, based on which one may define additional marker genes. This process can be conducted iteratively and monitored by other analyses to verify the biological significance of the findings.

Furthermore, newer developments of scRNA-Seq and similar technologies have enabled multiplexing of diverse sources of data such cellular localization, spatial organization and cell lineages. Supervised approaches, such as SCINA, could be advantageous in handling these challenging analytical tasks as the supervision process could effectively integrate other types of information (location, lineage, *etc)* naturally into the transcriptional profiles.

Overall, SCINA, a “signature-to-category” algorithm, addresses a critical research need that has been previously neglected. SCINA, when coupled with other “category-to-signature” methods, could greatly enhance the flexibility and power of researchers in the application of single cell profiling to derive exciting biological discoveries.

## METHODS

### The SCINA algorithm

SCINA is regarded as semi-supervised because the prior knowledge of signature genes is built into the unsupervised estimation process. This is different from supervised learning, where the goal is to minimize a loss function to approximate the known labels of instances. SCINA accepts a list of signature gene sets for a variety of cell types, and an expression matrix, which is assumed to have been pre-processed by the user with logarithmic transformation and/or any appropriate method of normalization if necessary. For each cell type, the signature can have one or more genes. The signature genes, by default, should be highly expressed in one particular cell type compared to all other cell types. Expression of genes that are characteristically lowly expressed in one cell type compared to the other cell types can be inverted so that the pseudo expression of this gene is high in that cell type. The SCINA model assumes that there is a bimodal distribution for each signature gene, with the higher mode corresponding to the cell type(s), in which this gene is designated as a signature, and the lower mode corresponding to all the other cell types. In the pool of cells being analyzed by SCINA, the group of cells without high expression of any of the signature genes will be designated as an “unknown” class of cells (default mode). However, SCINA also implements a switch that turns off the searching of “unknown” cells. The technical details of SCINA are specified in **Sup. File 1**.

### Simulation data generation

We simulated 30 lists of signature genes, with the size of each list being an integer drawn from a uniform distribution from 5 to 50. We assigned a number of cells to each simulated cell type, with the number generated according to the following distributions: (1) For the 30 (R) cell types, we assigned weights W(w_1_,w_2_…w_R_) generated randomly from a uniform distribution from 1 to 50. (2) For the R+1 signature (the ‘unknown’ cell type) we assigned the weight w_R+1_ generated randomly from a uniform distribution from 5 to 15. (3) For cell type *n*, the number of cells is 2000\**w_n_*/Σ(w_1_,w_2_…w_R_+1). For each gene, in the set of cells with this gene being designated as a signature, the gene expression was simulated from a normal distribution, whose mean was drawn from a uniform distribution between 0 and 3, and standard deviation drawn from another uniform distribution between 10^−5^ and 3. For the expression of this gene in other cell types, the expression was drawn from a normal distribution with mean drawn from a uniform distribution between 0 and 0.001, and standard deviation drawn from a uniform distribution between 10^−5^ and 3. Expression of genes in the ‘unknown’ class of cells was also simulated from the second distribution.

Next we challenged the SCINA algorithm by adding noise to the signatures with three methods: (1) When constructing the expression matrix, only R*-n* signatures were really used, while the other *n* signatures were not used and became “additional” gene signatures for non-existing cell types; (2) We simulated the expression matrix with all signatures but removed signatures of *n* cell types from the full list from the input of SCINA; (3) For each of *n* cell types, we added noise genes randomly selected from non-signature genes into that signature to double the size of the signature. We increased *n* from 1 to 29 in 29 tests for all three scenario. (4) Simulation of dropouts. In the expression simulation of each signature gene for each cell, we randomly performed dropout (expression is from the lower mode, not the higher mode) with a chance of n × 3.33% (n is from 1 to 10), depending on the empirical drop-out rates of single cell RNA-Seq data (typically 5% to 15%) (Grün et al., 2014).

### *Stk4* KO mouse related experiments

The generation of *Stk4* KO mice and the flow cytometry experiments follow the protocols that we have described elsewhere (Wang et al., 2015). For the Complete Blood Count (CBC) analysis, blood was collected into EDTA-coated MiniCollect blood collection tubes (Greiner Bio-One, Kremsmünster, Austria, catalog #VG-450474) via submandibular vein puncture from unanesthetized mice. 40 uL of blood from each mouse were aliquoted and analyzed by a Hemavet 950FS (Drew Scientific, Oxford, CT). Using an electrical impedance and focused flow cell system, the Hemavet quantified the total number of white blood cells, eosinophils, neutrophils, basophils, and lymphocytes.

### UTSW FHD/FHD-like patient cohort

RNA-seq and exome-seq samples of RCC patients from our UTSW Kidney Cancer Program (KCP) database and those mentioned in Durinck *et al* (*Durinck et al., 2015*) were pre-processed and analyzed. The FHD and FHD-like RCCs from 9 patients (XP108, XP515, UTSW85, UTSW86, UTSW87, XP878, XP552, UTSW88, and XP790) were reviewed by two expert genitourinary pathologists at UTSW (P.K.) and Memorial Sloan Kettering Cancer Center (V.R.). DNA and RNA were extracted and sequenced from their frozen tissue (5 patients) or FFPE blocks (4 patients) as available. Human subject study was performed in accordance with the protocol approved by the Institutional Review Board of the University of Texas Southwestern Medical Center. Approval numbers: 012011-190 and STU-22013-052.

### Genomics analysis pipelines

RNA-seq libraries were prepared using the TruSeq RNA Sample Preparation kit (Illumina). The libraries were multiplexed three per lane and sequenced on the HiSeq 2500 platform to obtain, on average, ~100 million paired-end (2 × 75-bp) reads per sample. Tophat2 with parameters “--num-threads 12 -g 10 -library-type fr-unstranded” was used to align the RNA-seq reads to the human genome (hg19). FeatureCounts(Liao et al., 2014) with parameters “-t exon -g gene_id -s 0 -T 12 --largestOverlap -- minOverlap 3 -M --fraction --ignoreDup -p -P -B -C” was used to count gene expression levels.

Exome capture was performed using the Agilent SureSelect Human All Exon kit (50Mb). Exome capture libraries were sequenced on the HiSeq 2500 platform (Illumina) to generate 2 × 75-bp paired-end data. Quality of exome-seq data was examined by NGS-QC-Toolkit (Patel and Jain, 2012). Exome-seq reads were aligned to the hg19 genome by BWA-MEM (Li and Durbin, 2009). Picard was used to add read group information and mark PCR duplicates. A GATK toolkit(DePristo et al., 2011; McKenna et al., 2010; Van der Auwera et al., 2013) was used to perform base quality score recalibration and local realignment around Indels. GATK HaplotypeCaller with SNP and Indel recalibration, MuTect (Cibulskis et al., 2013), and VarScan (Koboldt et al., 2012) were used to call SNPs and Indels. Annovar was used to annotate SNPs and Indels (Wang et al., 2010). Only non-silent missense mutations that were predicted to be deleterious by either SIFT or Polyphen2 or loss-of-function mutations were kept. Somatic mutations and germline mutations were annotated according to the mutation allele frequencies in the tumor and normal samples. Mutations that had a background mutation frequency >1% in any of ExAC, Esp6500, 1000 Genome, HRC, or Kaviar were eliminated.

### Statistical analyses

All computations and statistical analyses were carried out in the R computing environment. K-means clustering was carried out using the CellRanger software suite (Zheng et al., 2017) (version 2.2.0). Reanalysis functions provided by the package, including run_pca and run_kmeans_clustering, were applied with default parameters, except for the enforcing of two clusters for the k-means clustering. For running t-SNE, we used the Rtsne R package (version 0.13). We set dims=2 and theta=1, and used default values for all other parameters. Seurat analysis was performed with the Seurat R toolkit (version 2.3.4), the functions and parameters setting followed the tutorial provided on the Seurat website (https://satijalab.org/seurat/pbmc3k_tutorial.html). SINCERA analysis was applied with the SINCERA R package (version 0.99.0), following the vignette from the Xu Lab (https://github.com/xu-lab/SINCERA/blob/master/demo/mouselung.e16.5.R). PhenoGraph analysis was applied with the Rphenograph R package (version 0.99.1). The parameters were adjusted according to the dataset sizes to achieve the best performances. FACS/CyTOF data were downloaded or generated as .fcs files and converted to numerical matrices with the R package cytofkit (Chen et al., 2016) (version 1.6.5). Gene ontology analyses were carried out using the GOrilla server (Eden et al., 2009, 2007) (http://cbl-gorilla.cs.technion.ac.il/). Principal Component Analysis of RCC gene expression data was conducted with scaling. All statistical tests are two-tailed. For all boxplots appearing in this study, box boundaries represent interquartile ranges, whiskers extend to the most extreme data point which is no more than 1.5 times the interquartile range, and the line in the middle of the box represents the median.

## CODE ACCESS

The SCINA R package is available at https://github.com/jcao89757/SCINA and https://cran.r-project.org/web/packages/SCINA. The SCINA web server is available at http://lce.biohpc.swmed.edu/scina. All gene signatures used in this study are provided publicly on the SCINA web server.

## DATA ACCESS

UTSW RCC patients were asked if they would specifically consent to placement of their raw genomic data in a protected publicly accessible database. The RNA-Seq and exome-seq data of 5 consented UTSW FHD/FHD-like patients can be downloaded from the European Genome-phenome Archive with accession number EGAS00001002646 through controlled access. Other available KCP patient data can be accessed in EGAS00001000926.

## FUNDING

This study was supported by the National Institutes of Health (NIH) [R03 ES026397-01/TW, ZZ; R01 CA175754/JB; SPORE 1P50CA19651601/JB, TW, PK; CCSG 5P30CA142543/TW], Center for Translational Medicine of UT Southwestern [SPG2016-018/TW], and Cancer Prevention and Research Institute of Texas [CPRIT RP150596/DL]. This work was also partially supported by fundraising efforts orchestrated by the KCP Patient Council and the Kidney Cancer Coalition.

## ACKNOWLEDGEMENTS

We would like to thank Jessie Norris and Dr. Richie Xu for their helpful comments on the writing of the paper and Drs. Steffen Durinck and Kenian Chen for their helpful input on the scientific content of this work. We would also like to thank Dr. Bruce Beutler for generating the Stk4-related mouse data.

## AUTHOR CONTRIBUTIONS

Z.Z. and S.W. contributed to the computational analyses. D.L. designed the SCINA web server. X.Z., J.H.C., and E.M. contributed to the Stk4-related experiments. Y.M., E.S., Z.M., and S.S. contributed to the sequencing experiments of the FHD/FHDL samples. P.K. contributed to the pathological analyses of the FHD/FHDL samples. T.W. contributed to the overall supervision of the project. X.W., G.H., J.B., and T.W. contributed to study design and manuscript writing.

---

**Sup. Fig. 1.**
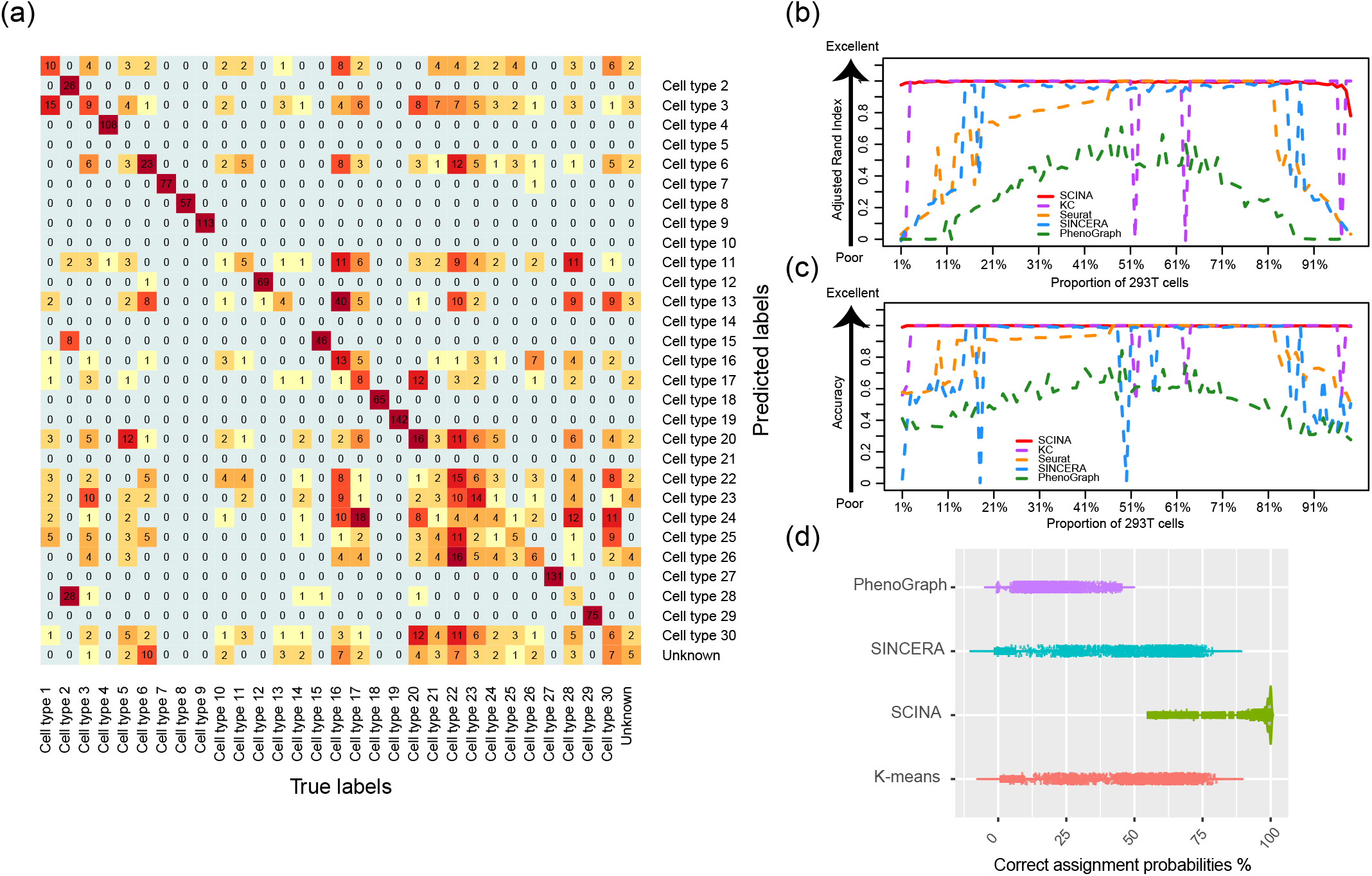
Comparison of SCINA and other algorithms on simulation data. (a) KC was performed on the same simulated dataset as in **Fig. 2a**. The heatmap showed the overlap between the simulated cell types and the detected cell types by KC. Each cluster detected by KC was assigned to the simulated cell type with the largest overlap with that cluster. (b) SCINA, KC, Seurat, SINCERA and PhenoGraph were used to identify Jurkat T cells and HEK293 cells. Accuracy is measured by Adjusted Random Index (ARI). (c) The same analyses evaluated by percentage of cells correctly assigned (ACC). (d) Violin plot showing probabilities of all simulated cells being assigned to their ‘true’ cell types. Analysis were performed on 100 datasets, each with 2000 cells simulated with the same approaches as in **Fig. 2a**.

**Sup. Fig. 2.**
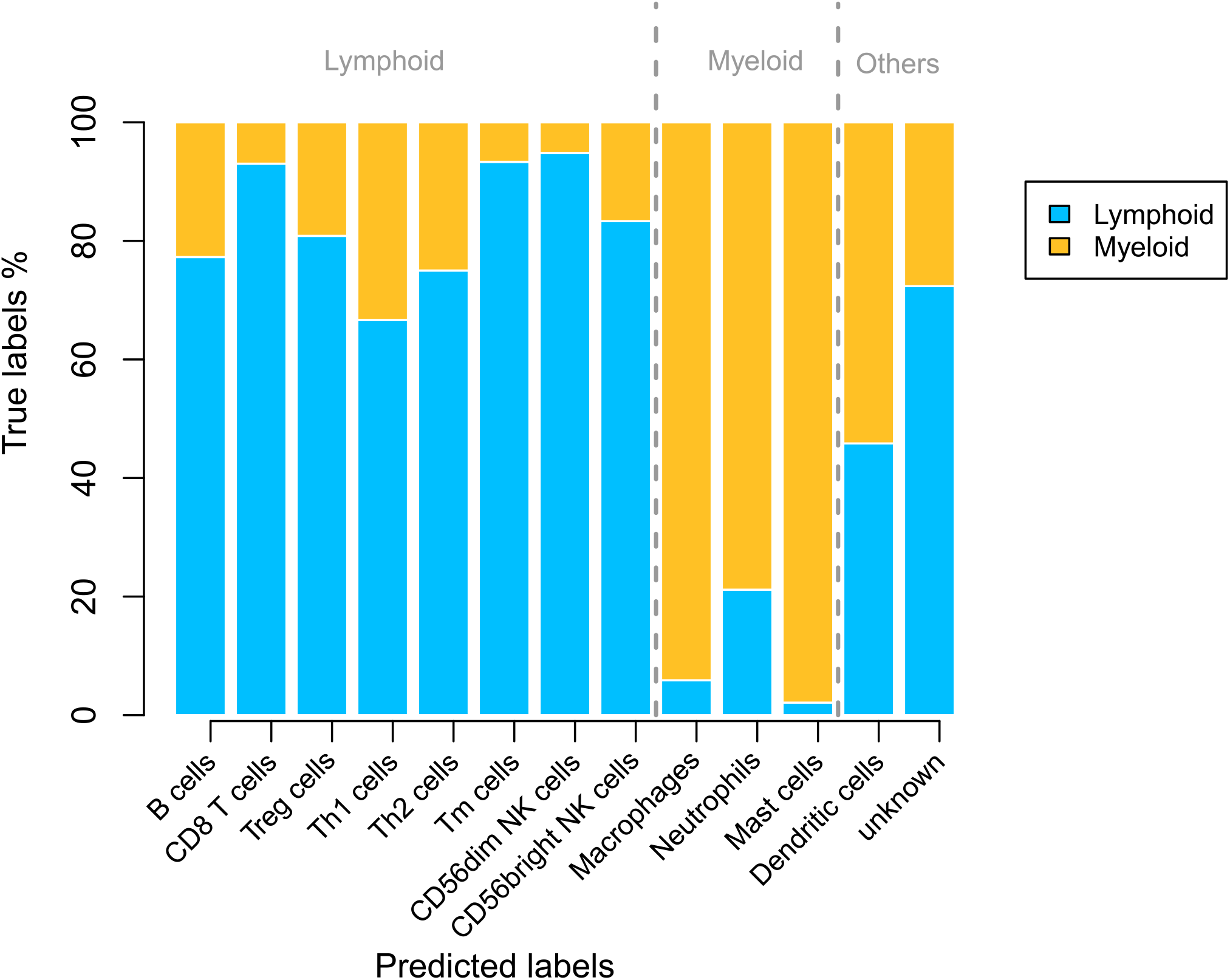
SCINA identified cell types of the CD45+ single cells enriched from the RCC tumor microenvironment. Bars are colored differently according to the gold standard of the lymphoid and myeloid pool labels.

## Notes

**Conflict of interest:** The authors declare no conflict of interest.

