## Supplement. File 1 for "SCINA: Semi-Supervised Analysis of Single Cells *in silico*"

### The Semi-supervised Category Identification and Assignment (SCINA) algorithm

Notations:

Genes: $i = 1...I$

Cells: $j = 1...J$

Single cell profiling data (normalized and log transformed as needed): $E_{ij}$

Cell types to be assigned: $r = 1...R$ . The cell types are mutually exclusive.

Gene signatures for each cell type: $S_r$ . Overlap between signatures is discouraged

True identity of each profiled cell: $z_j = 0,1,...R$

Goal:

Determine the cell type of each profiled cell, j, based on signature genes. By default, activation of gene signatures are assumed. Expression of signature genes that are characteristically lowly expressed in one cell type compared to the other cell types can be inverted so that the pseudo expression of this gene is high in that cell type.

Each cell, j, belongs to either one of the R cell types or other cells types (novel unknown cell types) ($z_j = 0$). Each type of cells, r, is marked by activation of the signature genes, $S_r$ . It can be assumed the unknown cell type does not have activation of any gene signature.

Modeling assumptions

(1) For each cell type's signature genes, assume the expression across all sequenced cells follow a bi-modal distribution:

$$(\overrightarrow{e_{S_r,j}} \mid z_j = r) \sim N(\overrightarrow{\mu_{r,1}}, \Sigma_1^r)$$
$$(\overrightarrow{e_{S_r,j}} \mid z_j \neq r) \sim N(\overrightarrow{\mu_{r,2}}, \Sigma_2^r)$$

$\overrightarrow{\mu_{r,1}}$ and $\overrightarrow{\mu_{r,2}}$ are vectors of means, and satisfy $\overrightarrow{\mu_{r,1}} > \overrightarrow{\mu_{r,2}}$ for each element of the vector

The covariance $\Sigma_1^r = \Sigma_2^r$ is a diagonal matrix

(2) Therefore, for each cell, j, to be analyzed:

If $z_j = s, s > 0$ , we have $\overrightarrow{e_{S_s,j}} \sim N(\overrightarrow{\mu_{s,1}}, \Sigma_1^s)$ and $\overrightarrow{e_{S_r,j}} \sim N(\overrightarrow{\mu_{r,2}}, \Sigma_2^s)$ for all $r \neq s, r = 1...R$

If $z_j = 0$ , we have $\overrightarrow{e_{S_r,j}} \sim N(\overrightarrow{\mu_{r,2}}, \Sigma_2^r)$ for all $r = 1...R$

(3) The idea can be conceptualized as in the following figure

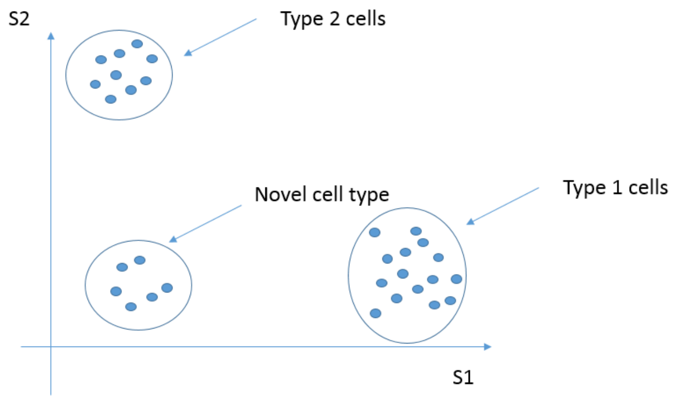


<u>E-M algorithm</u>

(1) Likelihood function

Let $P(z_j = r) = \tau_r, r = 0 \ldots R, \sum_{r=0 \ldots R} \tau_r = 1$

Parameters to be estimated: $\Theta = (\tau_r, \overrightarrow{\mu_{r,1}}, \overrightarrow{\mu_{r,2}}, \Sigma_1^r, \Sigma_2^r)$ , for all $r = 1 \ldots R$

$$L(\Theta; E) = \prod_{j=1 \ldots J} \{\tau_0 \prod_{r=1 \ldots R} f(\overrightarrow{e_{S_r,j}}; \overrightarrow{\mu_{r,2}}, \Sigma_2^r) + \sum_{s=1 \ldots R} [\tau_s f(\overrightarrow{e_{S_s,j}}; \overrightarrow{\mu_{s,1}}, \Sigma_1^s) \prod_{r=1 \ldots R, r \neq s} f(\overrightarrow{e_{S_r,j}}; \overrightarrow{\mu_{r,2}}, \Sigma_2^r)]\}$$

(2) E step

$$P^t(z_j = 0 \mid E; \Theta^t) = \frac{\tau_0^t \prod_{r=1 \ldots R} f(\overrightarrow{e_{S_r,j}}; \overrightarrow{\mu_{r,2}^t}, \Sigma_2^{r,t})}{\tau_0^t \prod_{r=1 \ldots R} f(\overrightarrow{e_{S_r,j}}; \overrightarrow{\mu_{r,2}^t}, \Sigma_2^{r,t}) + \sum_{s=1 \ldots R} [\tau_s^t f(\overrightarrow{e_{S_s,j}}; \overrightarrow{\mu_{s,1}^t}, \Sigma_1^{s,t}) \prod_{r=1 \ldots R, r \neq s} f(\overrightarrow{e_{S_r,j}}; \overrightarrow{\mu_{r,2}^t}, \Sigma_2^{r,t})]}$$

or

$$= \frac{\tau_0^t}{\tau_0^t + \sum_{s=1 \ldots R} [\tau_s^t \frac{f(\overrightarrow{e_{S_s,j}}; \overrightarrow{\mu_{s,1}^t}, \Sigma_1^{s,t})}{f(\overrightarrow{e_{S_s,j}}; \overrightarrow{\mu_{s,2}^t}, \Sigma_2^{s,t})}]}$$

$$P^t(z_j = s_0, s_0 > 0 \mid E; \Theta^t) = \frac{\tau_{s_0}^t f(\overrightarrow{e_{S_{s_0},j}}; \overrightarrow{\mu_{s_0,1}^t}, \Sigma_1^{s_0,t}) \prod_{r=1 \ldots R, r \neq s_0} f(\overrightarrow{e_{S_r,j}}; \overrightarrow{\mu_{r,2}^t}, \Sigma_2^{r,t})}{\tau_0^t \prod_{r=1 \ldots R} f(\overrightarrow{e_{S_r,j}}; \overrightarrow{\mu_{r,2}^t}, \Sigma_2^{r,t}) + \sum_{s=1 \ldots R} [\tau_s^t f(\overrightarrow{e_{S_s,j}}; \overrightarrow{\mu_{s,1}^t}, \Sigma_1^{s,t}) \prod_{r=1 \ldots R, r \neq s} f(\overrightarrow{e_{S_r,j}}; \overrightarrow{\mu_{r,2}^t}, \Sigma_2^{r,t})]}$$

$$= \frac{\tau_{s_0}^t \frac{f(\overrightarrow{e_{S_{s_0},j}}; \overrightarrow{\mu_{s_0,1}^t}, \Sigma_1^{s_0,t})}{f(\overrightarrow{e_{S_{s_0},j}}; \overrightarrow{\mu_{s_0,2}^t}, \Sigma_2^{s_0,t})}}{\tau_0^t + \sum_{s=1 \ldots R} [\tau_s^t \frac{f(\overrightarrow{e_{S_s,j}}; \overrightarrow{\mu_{s,1}^t}, \Sigma_1^{s,t})}{f(\overrightarrow{e_{S_s,j}}; \overrightarrow{\mu_{s,2}^t}, \Sigma_2^{s,t})}]}$$

$$\frac{f(\vec{e};\overrightarrow{\mu_1},\Sigma_1)}{f(\vec{e};\overrightarrow{\mu_2},\Sigma_2)} = \frac{|\Sigma_2|^{0.5}}{|\Sigma_1|^{0.5}} e^{-\frac{1}{2}(\vec{e}-\overrightarrow{\mu_1})^T \Sigma_1^{-1}(\vec{e}-\overrightarrow{\mu_1})+\frac{1}{2}(\vec{e}-\overrightarrow{\mu_2})^T \Sigma_2^{-1}(\vec{e}-\overrightarrow{\mu_2})}$$

(3) M step

Updating $\tau_r^{t+1} = \dfrac{\sum_{j=1...J} P^t(z_j = r \mid E;\Theta^t)}{J}$

Define

$$Q(\Theta \mid \Theta^t) = \sum_{j=1}^{J} \begin{array}{l} \{P^t(z_j = 0 \mid E;\Theta^t)[\log(\tau_0) + \sum_{r=1}^{R} \log f(\overrightarrow{e_{S_r,j}};\overrightarrow{\mu_{r,2}},\Sigma_2^r)] + \\ \sum_{s=1}^{R} P^t(z_j = s \mid E;\Theta^t)[\log(\tau_s) + \log f(\overrightarrow{e_{S_s,j}};\overrightarrow{\mu_{s,1}},\Sigma_1^s) + \sum_{\substack{r=1...R \\ r\neq s}} \log f(\overrightarrow{e_{S_r,j}};\overrightarrow{\mu_{r,2}},\Sigma_2^r)]\} \end{array}$$

Then

$$(\overrightarrow{\mu_{s,1}^{t+1}},\overrightarrow{\mu_{s,2}^{t+1}},\Sigma_1^{s,t+1},\Sigma_2^{s,t+1}) = \underset{\overrightarrow{\mu_{s,1}},\overrightarrow{\mu_{s,2}},\Sigma_1^s,\Sigma_2^s}{\arg\max} \sum_{j=1}^{J} \begin{array}{l} \{P^t(z_j = s \mid E;\Theta^t)\log f(\overrightarrow{e_{S_s,j}};\overrightarrow{\mu_{s,1}},\Sigma_1^s) + \\ \sum_{\substack{r=0...R \\ r\neq s}} P^t(z_j = r \mid E;\Theta^t)\log f(\overrightarrow{e_{S_s,j}};\overrightarrow{\mu_{s,2}},\Sigma_2^s)\} \end{array}$$

$$= \underset{\overrightarrow{\mu_{s,1}},\overrightarrow{\mu_{s,2}},\Sigma_1^s,\Sigma_2^s}{\arg\max} \sum_{j=1}^{J} \begin{array}{l} \{P^t(z_j = s \mid E;\Theta^t)[-\frac{1}{2}\log|\Sigma_1^s| - \frac{1}{2}(\overrightarrow{e_{S_s,j}} - \overrightarrow{\mu_{s,1}})^T \Sigma_1^{s,-1}(\overrightarrow{e_{S_s,j}} - \overrightarrow{\mu_{s,1}})] + \\ \sum_{\substack{r=0...R \\ r\neq s}} P^t(z_j = r \mid E;\Theta^t)[-\frac{1}{2}\log|\Sigma_2^s| - \frac{1}{2}(\overrightarrow{e_{S_s,j}} - \overrightarrow{\mu_{s,2}})^T \Sigma_2^{s,-1}(\overrightarrow{e_{S_s,j}} - \overrightarrow{\mu_{s,2}})]\} \end{array}$$

Updating

$$\overrightarrow{\mu_{s,1}^{t+1}} = \frac{\sum_{j=1}^{J} P^t(z_j = s \mid E;\Theta^t)\overrightarrow{e_{S_s,j}}}{\sum_{j=1}^{J} P^t(z_j = s \mid E;\Theta^t)}$$

$$\overrightarrow{\mu_{s,2}^{t+1}} = \frac{\sum_{j=1}^{J}\sum_{\substack{r=0...R \\ r\neq s}} P^t(z_j = r \mid E;\Theta^t)\overrightarrow{e_{S_s,j}}}{\sum_{j=1}^{J}\sum_{\substack{r=0...R \\ r\neq s}} P^t(z_j = r \mid E;\Theta^t)} = \frac{\sum_{j=1}^{J}[1 - P^t(z_j = s \mid E;\Theta^t)]\overrightarrow{e_{S_s,j}}}{\sum_{j=1}^{J}[1 - P^t(z_j = s \mid E;\Theta^t)]}$$

If calculated in this way and any gene i in the gene set $S_r$ satisfy $\mu_{s,1}^{t+1}(i) < \mu_{s,2}^{t+1}(i)$ , then

$$\mu_{s,1}^{t+1}(i) = \mu_{s,2}^{t+1}(i) = \frac{\sum_{j=1...J} e_{i,j}}{J}$$

Updating covariance

$$D_{s,1}^{t+1} = diag((\overrightarrow{e_{S_s,j}} - \overrightarrow{\mu_{s,1}^{t+1}}) \circ (\overrightarrow{e_{S_s,j}} - \overrightarrow{\mu_{s,1}^{t+1}}))$$

$$D_{s,2}^{t+1} = diag((\overrightarrow{e_{S_s,j}} - \overrightarrow{\mu_{s,2}^{t+1}}) \circ (\overrightarrow{e_{S_s,j}} - \overrightarrow{\mu_{s,2}^{t+1}}))$$

$$\Sigma_{s,1}^{t+1} = \Sigma_{s,2}^{t+1} = \frac{\sum_{j=1}^{J}[1 - P^t(z_j = s \mid E;\Theta^t)]D_{s,2}^{t+1} + P^t(z_j = s \mid E;\Theta^t)D_{s,1}^{t+1}}{J}$$

(4) Initial values

$$\tau_r^0 = \frac{1}{R+1}, r = 0...R$$

$\overrightarrow{\mu_{r,1}^0}, \overrightarrow{\mu_{r,2}^0}, r = 1...R$ assigned by quantiles of empirical distribution

$\Sigma^{r,0}, r = 1...R$ is a diagonal matrix with diagonal elements corresponding to variance of each gene multiplied by a constant

(5) Stop rules

Stabilization of assigned cell type labels

Overlap between gene signatures

This version of SCINA handles overlap between gene signatures in a heuristic manner. When there is overlap, the algorithm “virtually” creates multiple rows in the expression matrix for the overlapping genes. The number of virtual rows correspond to the number of times this gene appeared in the gene signature sets. Then in effect, this gene in each signature is assigned to each duplicate row, and the EM algorithm assumes these rows are independent of each other.

Con: The modeling assumption entails that each duplicate row has two modes, with the bigger mode corresponding to the expressional contribution from only one cell type and the smaller mode contains contributions from only non-expressing cells. This is violated with overlapping gene signatures. However, when these genes are only shared by a small subset of cell types compared to the whole population of cell types, the bias should be small.

Pro: this setup enables a clean separation between signatures of different cell types. Thus, SCENA benefits from a huge speedup due to the vectorized computation in R.
